# Molecular mechanisms of *Plasmodium* development in male and female *Anopheles* mosquitoes

**DOI:** 10.1101/2022.01.27.477980

**Authors:** Asako Haraguchi, Makoto Takano, Jun Hakozaki, Kazuhiko Nakayama, Sacré Nakamura, Yasunaga Yoshikawa, Shinya Fukumoto, Kodai Kusakisako, Hiromi Ikadai

## Abstract

Vector competence influences the ability of *Anopheles* mosquitoes to transmit *Plasmodium* parasites. The aim of this study was to determine the competence of male and female *Anopheles* mosquitoes to support the development of *Plasmodium* parasites. Male and female *A. stephensi* (STE2 strain) were infected with *in vitro*-cultured *P. berghei* (ANKA strain) ookinetes. We found that the number of oocysts produced was higher in males than in females. RNA-seq analysis of male and female mosquitoes injected with *P. berghei* ookinetes showed that predominantly genes of unknown function changed in expression levels in response to ookinete infection; however, further studies are required to elucidate their functions. Moreover, male mosquitoes were injected with *in vitro*-cultured *P. falciparum* (3D7 strain) gametocytes or zygotes. The development of *P. falciparum* in males was detected using nested polymerase chain reaction. We found the DNA content was higher on day 15 than on day 0, indicating that *P. falciparum* developed in the mosquito hemocoel. This study revealed promising new mechanisms underlying the interactions between *Plasmodium* and mosquitoes.

**Author Summary:** *Anopheles* mosquitoes transmit *Plasmodium* parasites that cause malaria disease. Vector competence is the ability of vectors to support pathogen development. The aim of this study was to determine the competence of male and female mosquitoes for *Plasmodium* parasites. Oocysts were formed in both male and female mosquitoes injected with ookinetes from *in vitro*-cultured *Plasmodium berghei*, which causes malaria in mice. The number of oocysts was higher in males, indicating that the competence in males was higher than that in females. RNA-seq analysis showed that the expression of genes of unknown function was highly variable in males and females. The genes defining the competence factors need further study, but the results indicate novel genes may be discovered. Furthermore, the development of *Plasmodium falciparum,* which causes the most severe malaria in humans, in male mosquitoes injected with *in vitro*-cultured gametocytes or zygotes was detected. As male mosquitoes do not suck blood, this method may allow us to experiment with *Plasmodium* more safely.

## Introduction

Malaria is a serious protozoan disease that infects 200 million and kills 400 000 people per year [1]. It is caused by *Plasmodium* parasites transmitted by *Anopheles* mosquitoes [2–4]. Soon after feeding on an infected vertebrate host, the gametocytes form gametes that, after fertilization, give rise to zygotes that, within approximately 24 h, develop into motile ookinetes. Ookinetes traverse the mosquito midgut epithelial cells and differentiate into early oocysts attached to the basal side of the epithelium, facing the hemocoel. Within 10–15 d, the oocysts mature, releasing thousands of sporozoites into the hemolymph. These sporozoites invade the salivary glands and are transmitted to the next host when mosquitoes feed on another vertebrate host.

Less than one-tenth of the gametocytes ingested by a female mosquito become ookinetes, and less than one-tenth of the ookinetes develop into oocysts [3]. Thus, *Plasmodium* parasites undergo high losses during their development in the mosquito gut. This is a vulnerable period in the life cycle of the parasites and a vulnerable stage for transmission control [3,5,6].

Differences in susceptibility to pathogens between male and female hosts have been reported in various animal species. Male mice are more susceptible to *Babesia microti* and *P. berghei* than females [7,8]. Vector competence is the ability of vectors to support pathogen development and is one of the factors that define the ability of vectors to transmit pathogens [2,9]. In the dipteran tsetse fly (*Glossina morsitans*) a blood-feeder and pathogen vector like the mosquito, both males and females feed on blood and transmit *Trypanosoma brucei.* Male and female vector competence differ, with male salivary glands being more conducive to *T. brucei* development than female salivary glands [10]. Male mosquitoes do not feed on blood, and there are no reports examining male mosquito competence.

It has been reported that *in vitro*-cultured *P. berghei* ookinetes microinjected into the body cavity (hemocoel) of female *Anopheline* mosquitoes develop into oocysts and form sporozoites [11,12]. Microinjection of *P. gallinaceum* ookinetes into the dipteran fruit fly *(Drosophila melanogaster)* also yields oocysts and sporozoites [13]. Fruit flies are not natural hosts, and oocysts are not formed via blood feeding or oral ingestion of ookinetes; however, oocysts and sporozoites are formed when microinjected into the hemocoel, which indicates that *P. gallinaceum* cannot traverse the fruit fly midgut wall [13]. Moreover, that macrophages in the fruit fly act as a *P. gallinaceum* exclusion mechanism was clarified by studying fruit flies microinjected with *P. gallinaceum* in detail [13]. Thus, microinjection can be used to study the *Plasmodium* exclusion system or vector competence. Recently, RNA-seq analysis has emerged as a simple and powerful tool for investigating the *Plasmodium* vector competence of mosquitoes [14,15].

The aim of this study was to compare the competence of male and female *Anopheles* mosquitoes for *Plasmodium* by determining oocyst numbers and conducting RNA-seq analysis to investigate underlying molecular mechanisms.

## Results

### Development of *P. berghei* in male mosquitoes injected with ookinetes

*P. berghei* oocysts were formed in male and female mosquitoes at 14 d post injection of 2000 ookinetes (Fig 1A). Oocysts were formed in the esophagus, dorsal diverticula, crop, foregut, midgut, Malpighian tubules, hindgut, rectum, and inner abdominal wall of both males and females. At the esophagus, dorsal diverticula, and crop, a median of 42.5 and 9.5 oocysts formed in males and females, respectively. At the foregut and midgut, a median of 9.5 and 6.0 oocytes was observed in males and females, respectively. In the Malpighian tube, hindgut, and rectum, a median of 18.5 and 5.0 oocytes was detected in males and females, respectively. Last, at the abdominal wall, a median of 60.5 and 30.0 oocytes formed in males and females, respectively (Fig 1B). Oocysts were formed mostly at the esophagus, dorsal diverticula, crop, and abdominal wall in both males and females. In addition, more oocysts were formed in males than in females at every site. At 28 d post injection, *P. berghei* sporozoites were associated with the salivary glands, dorsal diverticula, crop, wings, and legs in males, and with the salivary glands, wings, and legs in females (Fig 1C). Mice infection was confirmed at 4 to 6 d post-tail-vein administration of 2000–10000 sporozoites from the salivary glands and whole body of male- and female-injected mosquitoes (S1 Table). When mice were fed with injected females, infection was confirmed 3 d post-blood feeding, indicating that *P. berghei* sporozoites formed in both males and females were infectious to mice.

**Fig 1.**
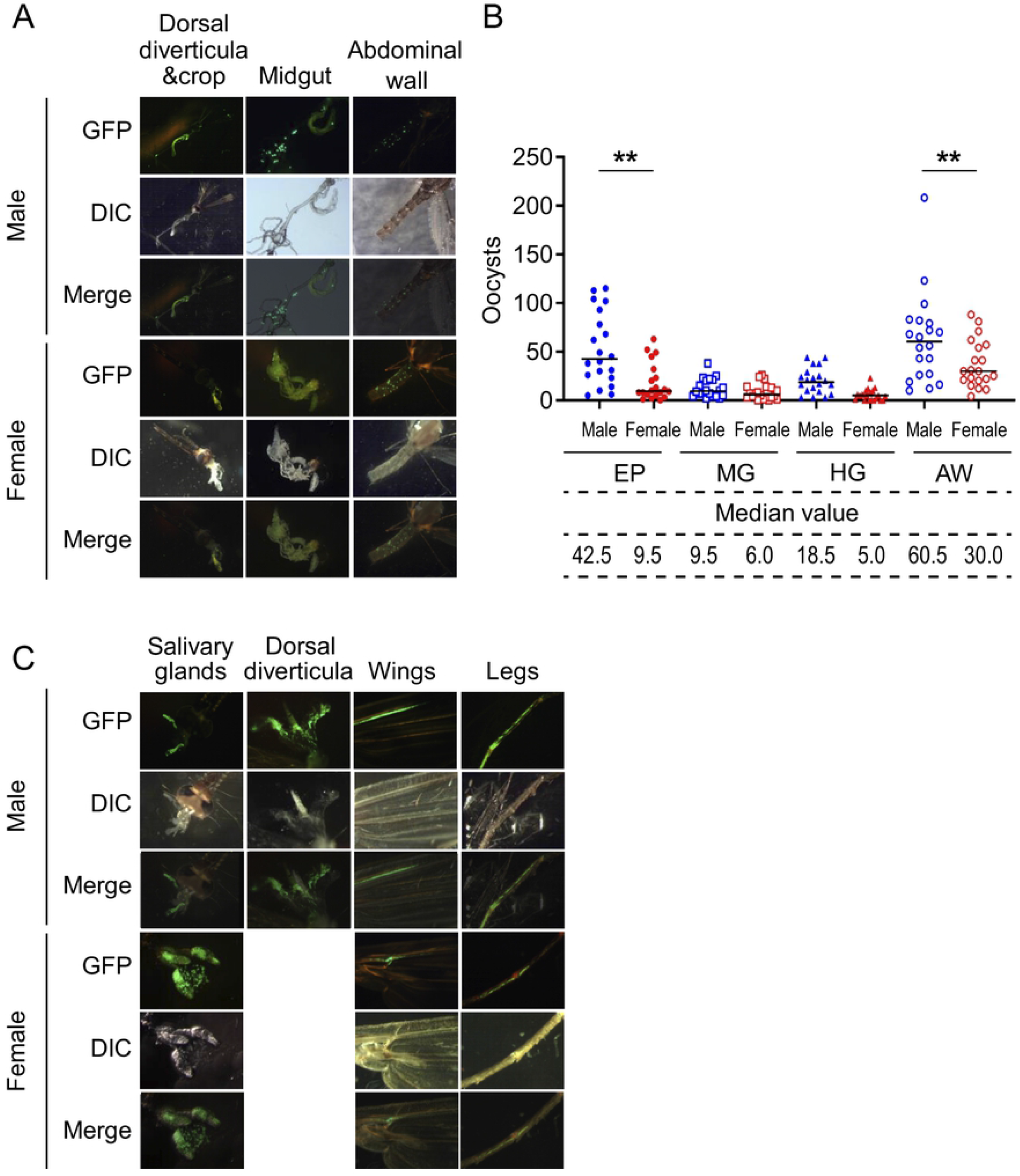
Oocyst and sporozoite formation in male and female mosquitoes injected with *Plasmodium berghei* ookinetes. (A) Oocysts in male and female mosquitoes viewed with a fluorescence microscope at 14 d after injection with 2000 GFP-expressing *P. berghei* ookinetes. DIC, differential interference contrast microscopy. (B) Number of oocysts formed at each site in males (n = 20) and females (n = 20). Mosquitoes were dissected at 14 d post injection. EP: esophagus, dorsal diverticula, and crop; MG: foregut and midgut; HG: Malpighian tube, hindgut, and rectum; AW: abdominal wall. The horizontal bar indicates the median value. Mann–Whitney test was used for comparison (\*\**p* < 0.01). Data are expressed as the pooled results of replicates. (D) Sporozoites in male and female mosquitoes under a fluorescence microscope at 28 d post injection with 2000 *P. berghei* GFP-expressing ookinetes.

### Higher oocyst number in male than in female mosquitoes

When 2000 ookinetes were injected into male and female mosquitoes, a median of 132 and 51 oocysts formed in males and females, respectively (Fig 2A). When 5000 ookinetes were injected, the median number of oocysts was 125 and 57 in males and females, respectively (Fig 2B). In both set-ups, the number of oocysts was significantly higher in males than in females (*p* < 0.01–0.05) and the prevalence was 100% in all mosquito groups.

**Fig 2.**
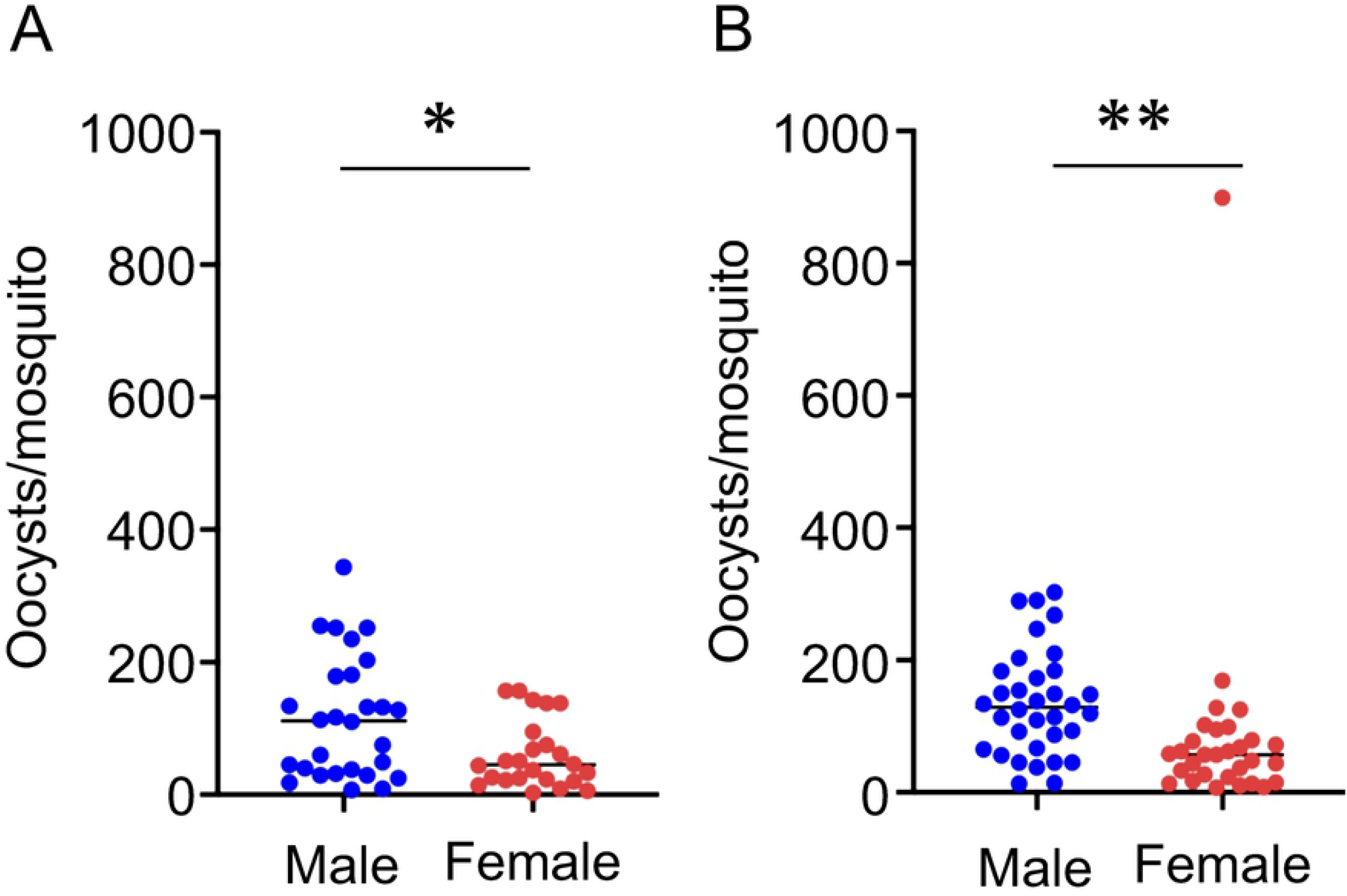
Oocyst numbers in male and female mosquitoes injected with *Plasmodium berghei* ookinetes. (A) Oocysts were counted at 14 d post injection of 2000 ookinetes in males (n = 28) and females (n = 24). (B) Oocysts were counted at 14 d post injection of 5000 ookinetes in males (n = 34) and females (n = 30). Horizontal bars indicate median values. Mann–Whitney test was used for comparison (\**p* < 0.05, \*\**p* < 0.01). Data are expressed as the pooled results of three replicates.

### RNA-seq analysis of male and female mosquitoes

To explore the genes that regulate *P. berghei* infection, we performed RNA-seq analysis of male and female mosquitoes at 24 h post injection, when ookinetes had differentiated into oocysts. More genes were differentially regulated in females than in males injected with 5 000 *P. berghei* ookinetes. In total, 140 genes were upregulated and 89 were downregulated in males, whereas 304 were upregulated and 167 were downregulated in females (fold change ≥|2|) (Fig 3A).

**Fig 3.**
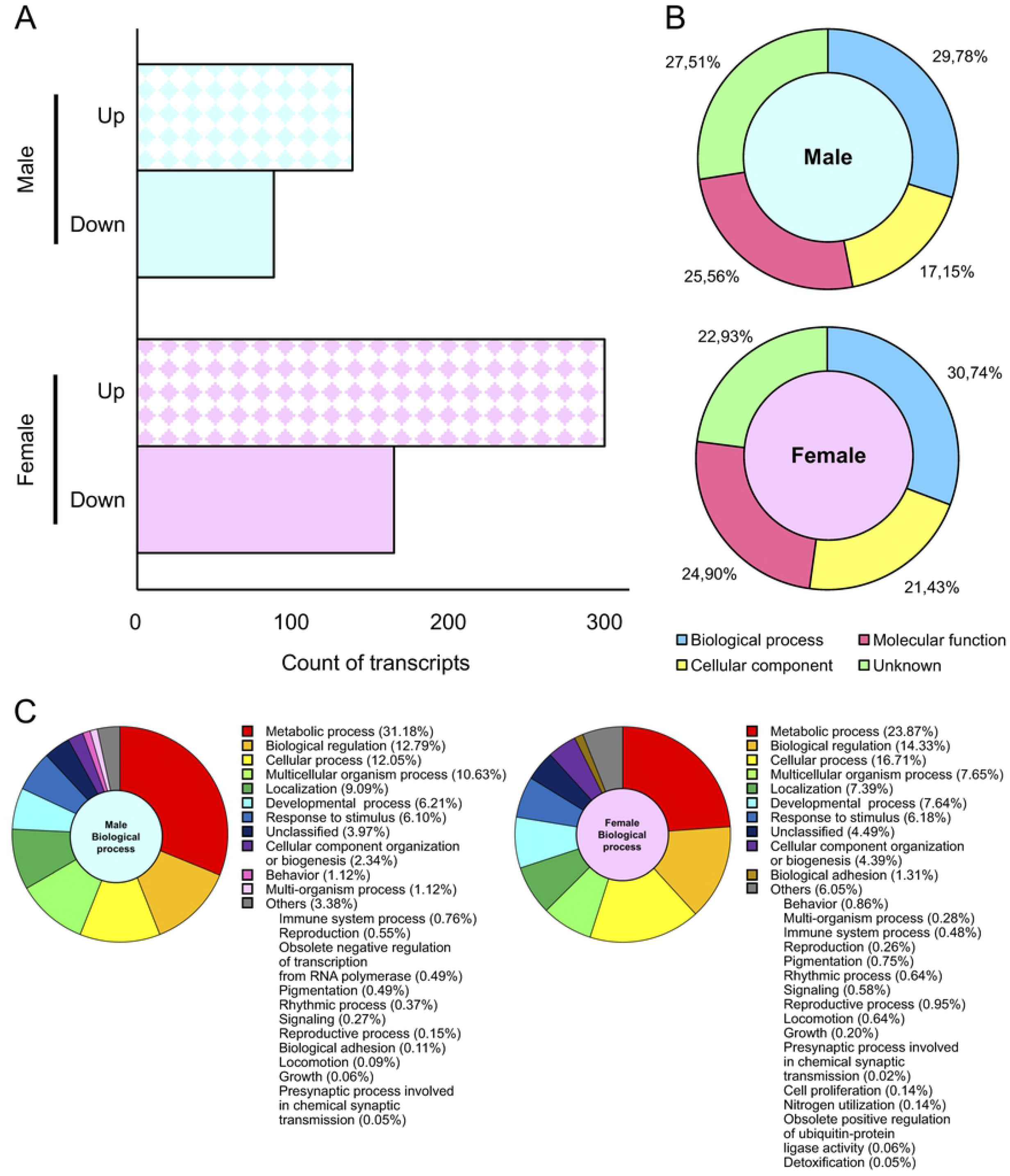
RNA-seq analysis of mosquitoes injected with *Plasmodium berghei* ookinetes. (A) Number of upregulated and downregulated genes in mosquitoes at 24 h post injection of 5000 *P. berghei* ookinetes (fold change ≥ |2|). Up: upregulated gene, Down: downregulated gene. (B) Percentage of variable genes annotated as biological process, cellular component, molecular function, or unknown function in gene ontology analysis. (C) Detailed functions of genes categorized as biological process.

Gene Ontology (GO) analysis showed there were 668 genes of known function and 193 genes with unknown functions. In males, 29.78% were classified as biological processes, 17.15% as cellular components, 25.56% as molecular functions, and 27.51% with unknown functions, whereas in females, the percentages were 30.74%, 21.43%, 24.90%, and 22.93%, respectively (Fig 3B). Among the biological processes, metabolic processes were the most common in both males and females, whereas immune system processes were observed at 0.76% in males and 0.48% in females (Fig 3C). The top 20 upregulated and downregulated genes in males and females are shown in Table 1. Most have unknown functions in both sexes.

**Table 1.**
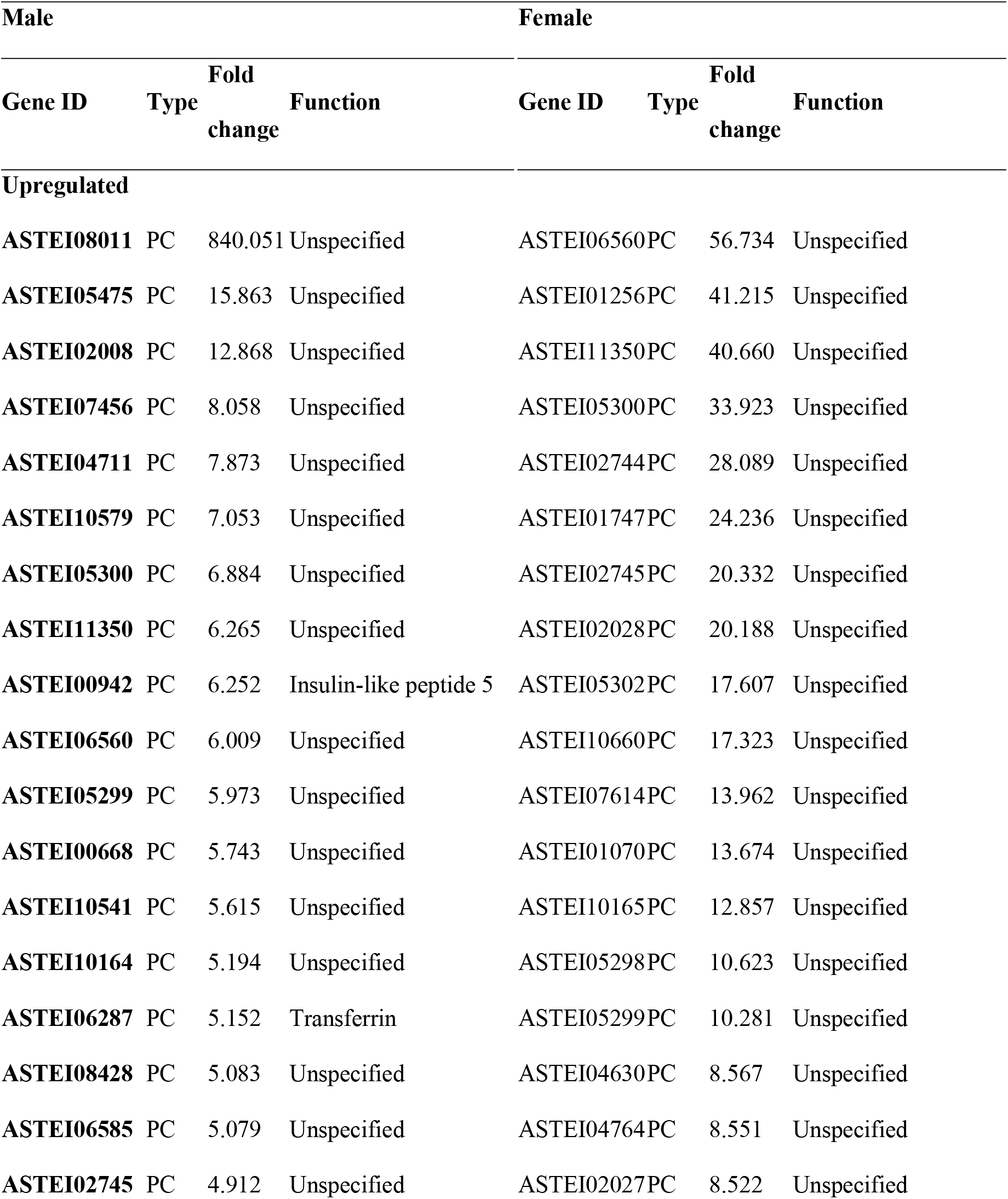
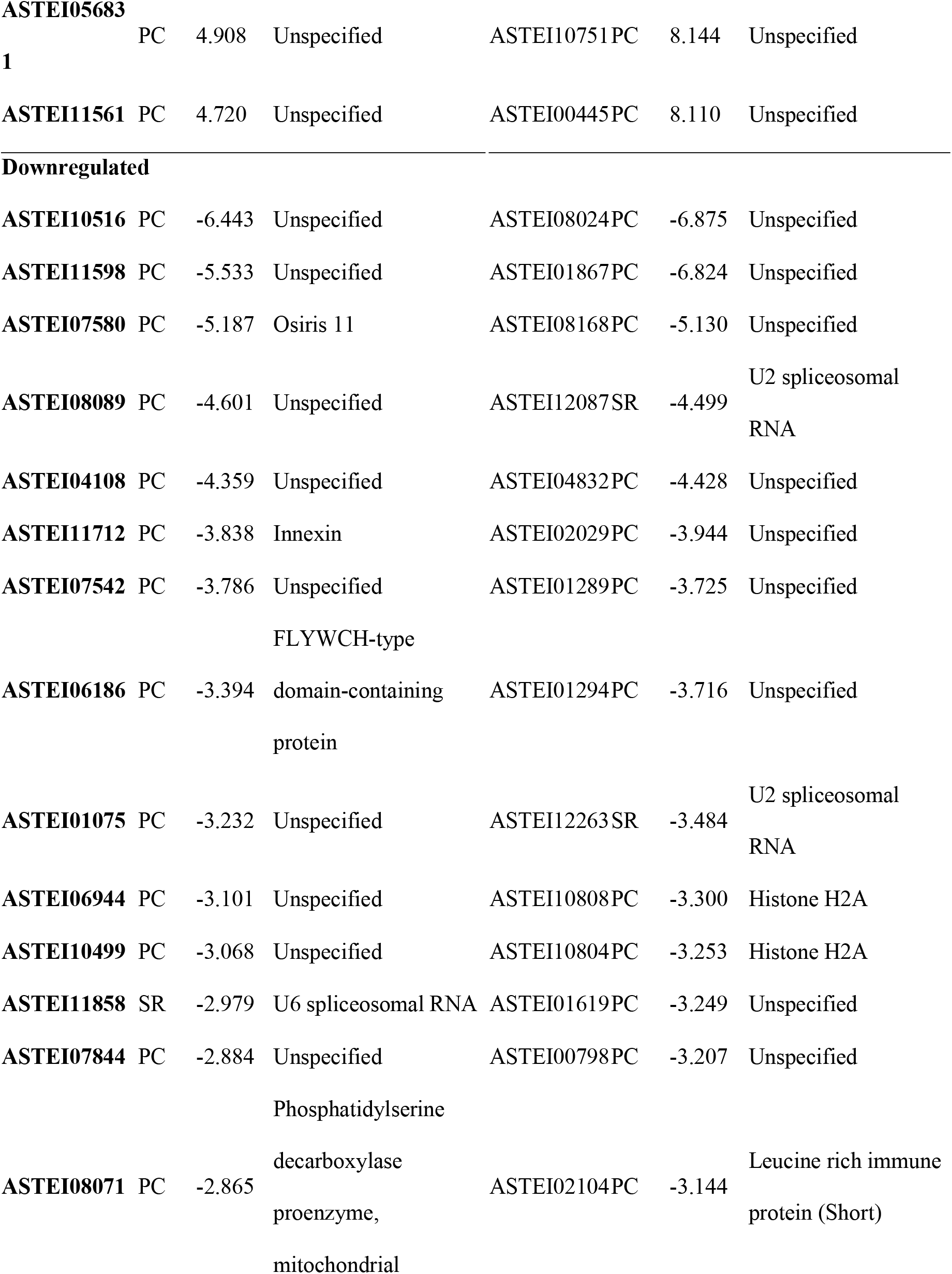
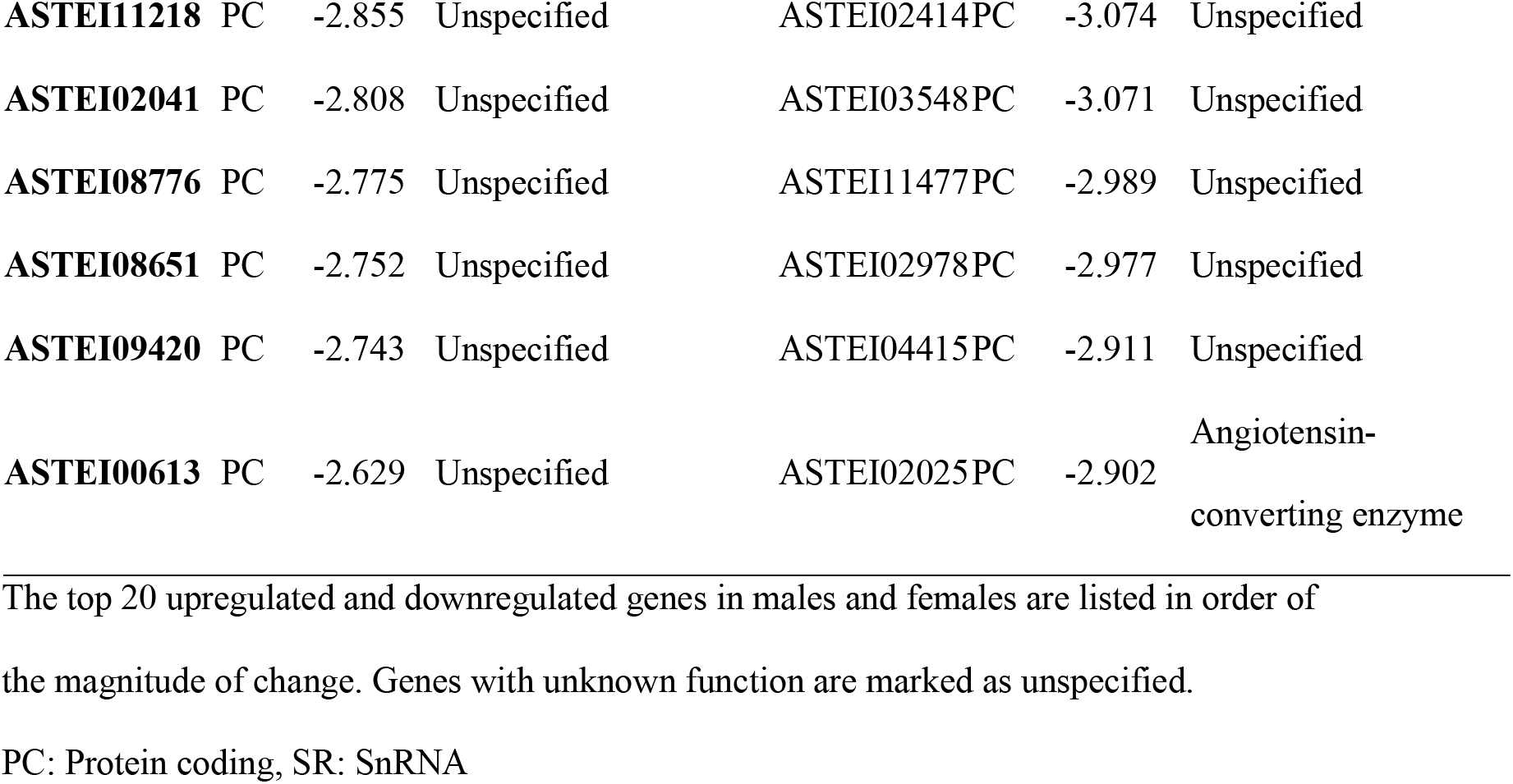
Genes with the most variable expression in male and female mosquitoes injected with *Plasmodium berghei* ookinetes.

### Detection of *P. falciparum* in male mosquitoes injected with gametocytes or zygotes

Male mosquitoes were injected with *P. falciparum* gametocytes or zygotes, and parasitic DNA in mosquitoes was measured on different days after injection. *P. falciparum* DNA was detected by nested polymerase chain reaction (PCR) up to 15 d post injection (Table 2). For 20 stage-V gametocytes injected into males, the detection rate of *P. falciparum* was 100% on day 0 and 30% on day 15. As for the 2, 1, and 1 stage-V gametocytes, the detection rates were 100.0%, 50.0%, and 81.3%, respectively, on day 0, and 30%, 0%, and 0%, respectively, on day 15. For 50 zygotes injected, the detection rate was 100% on days 0 to 11. Eight PCR-positive samples were sequenced and confirmed to be the *P. falciparum* 18S rRNA gene. In addition, the DNA content of *P. falciparum* in mosquitoes injected with 20 stage-V gametocytes was higher on day 15 than on day 0 (S1 Fig).

**Table 2.** Detection of *Plasmodium falciparum* in male mosquitoes injected with gametocytes or zygotes using nested polymerase chain reaction (PCR).

| Injected parasites<br>/mosquito | Positive rate of PCR [%]<br>(PCR-positive mosquitoes/total mosquitoes) |  |  |  |  |
| --- | --- | --- | --- | --- | --- |
|  | Days post injection |  |  |  |  |
|  | 0 | 1 | 11 | 12 | 15 |
| **20 gametocytes** | 100 (12/12) | 100 (2/2) | ND | ND | 30 (3/10) |
| **2 gametocytes** | 100 (6/6) | 0 (0/2) | ND | ND | 30 (3/10) |
| **1 gametocyte** | 50 (1/2) | 71 (5/7) | ND | ND | 0 (0/2) |
| **1 gametocyte** | 81 (13/16) | 40 (4/10) | 0 (0/1) | 10 (1/10) | 0 (0/17) |
| **50 zygotes** | 100 (13/13) | 100 (6/6) | 100 (1/1) | ND | ND |

Male mosquitoes were injected with the indicated number of parasites and at the indicated days, some mosquitoes were homogenized and assayed by PCR for the presence of *P. falciparum* 18S rRNA genes. Number of positive mosquitoes out of number analyzed is presented in parentheses. ND: No data

## Discussion

In this study, *P. berghei* oocysts formed throughout the body in male and female mosquitoes injected with ookinetes, particularly in the esophagus, dorsal diverticula, crop, and inner abdominal wall. In blood-feeding female mosquitoes, *P. berghei* ookinetes transform into oocysts by migrating through midgut epithelial cells to face the hemocoel [2–4]. It has been reported that *P. berghei* ookinetes injected into the female *A. gambiae* hemocoel can develop into oocysts anywhere in the hemocoel [11,12], consistent with the observations of this study. However, this study demonstrated that ookinetes can develop into oocysts in both male and female *A. stephensi* after injection. Many parasites migrate to specific tissues [16]; ookinetes injected into the hemocoel did not exhibit tissue specificity, suggesting that they migrated and adhered randomly to an available surface.

Furthermore, sporozoites were formed throughout the body at 28 d post injection and infected mice after administration into their tail vein or via mosquito blood feeding. Mosquitoes have an open circulatory system, and hemolymph from the dorsal vessels promotes circulation throughout the hemocoel [17,18]. Sporozoites move passively in the hemocoel following the hemolymph flow, attach to the salivary glands, and migrate into the salivary cavity [19]. They can be found attached to various tissues when mosquitoes are infected by blood feeding [5]. Thus, it is likely that sporozoites were transported throughout the mosquito body following the hemolymph flow and invaded the salivary glands.

Male mosquitoes exhibited more *P. berghei* oocysts than females at every site. The number of total oocysts in males was significantly higher than that in females (*p* < 0.01– 0.05), indicating that males exhibit greater competence than females. Moreover, injection of increased numbers of ookinetes did not increase the number of oocysts in both male and female mosquitoes. This suggests that a mosquito has a limited capacity to form an oocyst, or parasites can increase only up to a certain number; these results require further analysis. We attempted to investigate competence factors at the early oocyst stage using RNA-seq analysis at 24 h post injection. There were more gene expression changes in female than in male mosquitoes, those annotated to metabolic processes being the most common. Possibly, these genes participate in blocking early oocyst formation. Most up and downregulated genes in males and females have unknown functions, and differences in competence between males and females may be owing to these unknown-function genes. Vector competence in mosquitoes has been reported to be associated with many biological and environmental factors, such as physiological function, nutritional status, gut microbiota, and immune response [2,9]. Previous reports have compared strains or species of *Anopheles* mosquitoes with varying competence for *Plasmodium* parasites, indicating the involvement of the immune system and other factors [20–23]. However, in these studies, it was difficult to equalize genetic backgrounds and growth environments of mosquitoes, exclude these factors, and compare the competence factors. Therefore, the detailed molecular mechanisms that define vector competence remain unclear. As male and female mosquitoes of the same species and strain (e.g., *A. stephensi* [STE2 strain, MRA-128]), have the same genetic background and growth environment, a detailed examination of the genes revealed by the comparison of males and females in this study could unveil new facts concerning the molecular mechanisms of competence.

This study showed that *P. falciparum* infects male mosquitoes injected with gametocytes and zygotes, indicating that the development of gametocytes to ookinetes, which normally occurs in the midgut, is midgut-independent and occurs in the hemocoel. In blood-feeding mosquitoes, gametocytes differentiate into mature oocysts with sporozoites in 10–15 d [2–4]. The injection of *P. falciparum* gametocytes and zygotes showed infection at 15- and 11-d post injection, respectively. In addition, the quantity of *P. falciparum* DNA was greater on day 15 than on day 0 of gametocyte injection. There were differences among individuals, suggesting that mature oocysts with sporozoites formed in male mosquitoes. To confirm the development of a *P. falciparum* gametocyte, it is necessary to observe ookinetes, oocysts, and sporozoites in the injected mosquitoes. Currently, sporozoite vaccines to prevent the spread of *Plasmodium* parasites are being developed; however only one type of vaccine is currently in use, and more research on oocysts, which produce sporozoites internally, and sporozoites, which are infectious to humans and develop inside mosquitoes, is required to enable effective vaccine development [24,25]. A method for the *in vitro* culture of *P. falciparum* oocysts and sporozoites has been reported [26,27]; however, it is complicated and cumbersome, and the number of oocysts and sporozoites obtained is low. Injection of gametocytes into male mosquitoes may be a simpler way to develop oocysts and sporozoites, and as male mosquitoes are used, there is no risk of infection by blood feeding. This is expected to advance the study of the mosquito stages of these parasites.

## Conclusion

This study reveals that *P. berghei* ookinete injections into male and female mosquitoes, result in males having a higher competence than females. The differentially regulated genes in these males and females provide insight into the underlying molecular mechanisms involved in vector competence. In addition, *P. falciparum* infection succeeded in males when injected with gametocytes and zygotes. Hence, this study helps elucidate the factors involved in vector competence and access the mosquito stages of parasites more safely.

## Materials and Methods

### Mosquito rearing and microinjection of parasites

STE2 strain *A. stephensi* mosquitoes were reared in an insectary at 19 °C under a 14:10 h light:dark cycle with 10% (w/v) sucrose solution. A total of 69 nL of purified ookinete, gametocyte, and zygote suspensions was injected into the hemocoel of <10-day-old male and female mosquitoes using a Nanoject II automatic nanoliter injector (Drummond Scientific Company, Broomall, PA, USA) [11].

### *P. berghei* and ookinete culture

*P. berghei* (ANKA strain) that constitutively expresses green fluorescent protein (GFP) was maintained via serial passage in ICR mice (Charles River Laboratories Japan Inc., Yokohama, Japan) from frozen stock. Ookinete culture of *P. berghei* was performed as previously described [28,29]. Blood from infected mice was collected via cardiac puncture and placed in 10 volumes of ookinete medium containing RPMI medium 1640 (Thermo Fisher Scientific Inc., Waltham, MA, USA), 25 mM HEPES (Dojindo Laboratories, Kumamoto, Japan), 0.4 mM hypoxanthine (Sigma-Aldrich, St. Louis, MO, USA), 24 mM NaHCO_3_ (Fujifilm Wako Pure Chemical Corporation, Osaka, Japan), and 12.5 mg L^-1^ gentamicin reagent solution (Gibco) before incubation for 24 h at 19 °C.

### *P. falciparum* and culture of gametocytes and zygotes

*In vitro* culture of *P. falciparum* (3D7 strain) was performed according to the standard protocol [30] and maintained at 5% hematocrit in A^+^ human red blood cells (RBCs) on a complete medium at 37 °C under 5% CO_2_ and 5% O_2_. The complete medium comprised RPMI medium 1640, 25 mM HEPES, 0.1 mM hypoxanthine, 24–26 mM NaHCO_3_, 11 mM D(+)-glucose (Fujifilm Wako Pure Chemical Corporation), 12.5 mg L^-1^ gentamicin reagent solution, and 10% (v/v) heat-inactivated type A^+^ human plasma (Japan Red Cross Society, Tokyo, Japan). Gametocyte cultures were initiated from asexual parasites at 0.5–3.9% parasitemia and 6–10% hematocrit. The cultures were maintained for 14-27 d with daily medium changes without adding fresh RBCs. From day 1 to 5, the culture medium was supplemented with N-acetyl-D-glucosamine (Sigma-Aldrich) at a final concentration of 50 nM. Mature gametocytes were resuspended at 20% hematocrit in an exflagellation-inducing medium containing 10 mM Tris (pH 7.6, Fujifilm Wako Pure Chemical Corporation), 170 mM NaCl (Fujifilm Wako Pure Chemical Corporation), 25 mM NaHCO_3_, 10 mM D(+)- glucose, and 50–100 μM xanthurenic acid (Sigma-Aldrich) at 19 °C for 30 min for gamete maturation and zygote fertilization.

### Purification of parasites

*P. berghei* ookinetes and *P. falciparum* gametocytes and zygotes were purified using a MidiMACS separator system (LS Columns, Miltenyi Biotec, Bergisch Gladbach, Germany), as previously described [28,29]. Ookinetes, gametocytes, and zygotes were recovered by passing ookinete medium, incomplete medium comprising complete medium without 10% (v/v) heat-inactivated type A^+^ human plasma, and exflagellation-inducing medium, respectively.

### Examination of mosquitoes injected with *P. berghei*

Male and female mosquitoes were dissected at 14 d post injection, and oocyst localization, number, and prevalence were examined using an Eclipse E600 (Nikon, Tokyo, Japan) fluorescence microscope. At 28 d post injection, sporozoite localization was observed using a fluorescence microscope. Sporozoites from whole bodies or salivary glands of the injected males and females were administered into the tail vein of mice. Seven injected females were blood-fed on mice, and the infection in the mice was checked every day using a blood smear test.

### DNA extraction

Mosquitoes with *P. falciparum* gametocytes or zygotes from 0 to 15 d post injection were collected and stored at −80 °C. Whole mosquitoes were homogenized using a Micro Smash™ MS-100R (Tomy Seiko Co., Ltd., Tokyo, Japan) at 2500 rpm for 30 s. DNA was extracted using a ZymoBIOMICS Miniprep (Zymo Research Corporation, Irvine, CA, USA) according to the manufacturer’s protocol or using the phenol-chloroform isoamyl alcohol method. Briefly, a 500 μL genomic lysis buffer (Zymo Research Corporation) and 500 μL of phenol:chloroform:isoamyl alcohol (25:24:1) (Nippon Gene Co., Ltd., Tokyo, Japan) were added to extract the DNA, and then the DNA was precipitated using ethanol. The extracted DNA was dissolved with 10–50 μL of TE buffer and stored at −20 °C.

### Detection of *P. falciparum* using nested PCR

Prime STAR® HS (Premix) (Takara Bio Inc., Shiga, Japan) was used for PCR amplification, according to the manufacturer’s protocol, and the extracted DNA samples were used as template DNA. For *A. stephensi* S7 gene (loading control) amplification, PCR was conducted using primers AsRpS7-F (5’-CCTGGATAAGAACCAGCAGA-3’) and AsRpS7-R (5’-GGCCAGTCAGCTTCTTGTA-3’). The PCR conditions were as follows: 1 cycle of 95 °C for 3 min, followed by 35 cycles of 98 °C for 10 s, 55 °C for 5 s, and 72 °C for 18 s, and 1 cycle of 72 °C for 6 min. Distilled water as a negative control. Amplification of the *P. falciparum* 18S rRNA gene for detecting *P. falciparum* was conducted using nested PCR. The primers for the first PCR reaction were 18S rRNA-1F (5’-TTAATTTGACTCAACACGGGG-3’) and 18S rRNA-1R (5’-TATTGATAAAGATTACCTA-3’) [13]. The PCR conditions were as follows: 1 cycle of 95 °C for 3 min, followed by 40 cycles of 98 °C for 10 s, 55 °C for 15 s, and 72 °C for 33 s, and 1 cycle of 72 °C for 6 min. The primers for the second reaction were 18S rRNA-2F (5’-TAATAGCTCTTTCTTGAT-3’) and 18S rRNA-1R. The PCR conditions were as follows: 1 cycle of 95 °C for 3 min, followed by 35 cycles of 98 °C for 10 s, 55 °C for 15 s, and 72 °C for 30 s, and then 1 cycle of 72 °C for 6 min. Genomic DNA from *in vitro*-cultured *P. falciparum* was used as a positive control and the 7–11-d-old male mosquitoes as a negative control. Eight PCR-positive samples were sequenced using 18S rRNA-2F and 18S rRNA-1R primers (Fasmac Co., Ltd., Kanagawa, Japan).

### Real-time PCR

Among the PCR-positive mosquitoes injected with 20 stage-V gametocytes, DNA was quantified in three mosquitoes at 0 and 15 d post injection using KOD SYBR qPCR Mix (Toyobo Co., Ltd., Osaka, Japan), and 18S rRNA-3F (5’-GGATGGTGATGCATGGCCG-3’) and 18S rRNA-2R (5’-CAGTGTAGCACGCGTGCAG-3’) primers. The analysis was conducted using Applied Biosystems StepOnePlus (Thermo Fisher Scientific) with the following cycling conditions: 1 cycle at 98 °C for 3 min, followed by 40 cycles of 98 °C for 10 s, 60 °C for 10 s, and 68 °C for 30 s, and 1 cycle of 95 °C for 15 s, 60 °C for 1 min, and 99 °C for 15 s. *P. falciparum* DNA was quantified using a calibration curve, and each value was corrected by dividing by the quantity of template DNA used in the PCR reaction.

### RNA-seq analysis

Male and female mosquitoes at 24 h post injection with *P. berghei* ookinetes or with ookinete medium (negative control) were collected for RNA-seq analysis. Three mosquitoes were collected as one sample and stored at −80 °C. Total RNA was extracted using Nucleo Spin RNA Blood (Macherey-Nagel, Düren, Germany) according to the manufacturer’s protocol, delivered to Macrogen Japan Corp. (Tokyo, Japan) for library construction using the TruSeq Stranded mRNA LT Sample Prep Kit (Illumina, San Diego, CA, USA), and sequenced using the NovaSeq 6000 system (Illumina). FastQCv0.11.7 (http://www.bioinformatics.babraham.ac.uk/projects/fastqc/) was used to analyze the raw read quality, and statistical analysis was performed using fold change per comparison pair of mosquitoes injected with *P. berghei* ookinetes and with ookinete medium. GO analysis was performed using VectorBase (https://www.vectorbase.org/).

### Statistical analysis

All statistical analyses were performed using GraphPad Prism version 8.4.3 software (GraphPad Software Inc., San Diego, CA, USA). We determined the significance of oocyst intensity in injected mosquitoes using the Mann–Whitney test (*p* < 0.01–0.05) and the significance of DNA content for real-time PCR using the unpaired t-test (*p* < 0.05).

## Acknowledgments

The *Plasmodium falciparum* used in this study was kindly provided by Dr. Marcelo Jacobs-Lorena, John Hopkins University, MD, USA. The green fluorescent protein-expressing *Plasmodium berghei* ANKA strain was provided by Dr. Yuda, Mie University, Japan. The *Anopheles stephensi* (STE2 strain, MRA-128) was provided by BEI Resources. We are grateful to the Japanese Red Cross Society for providing the human A-type blood and A-type serum. We thank Dr. Marcelo Jacobs-Lorena for engaging in constructive discussions with us on particular aspects of this study. In addition, this study would not have been possible without the assistance of our laboratory support staff member Mai Tanaka. Finally, we would like to thank Editage (www.editage.com) for English language editing.

## Supporting Information

**S1 Fig. *Plasmodium falciparum* DNA content in male mosquitoes injected with gametocytes**. One mosquito injected with 20 stage-V gametocytes was used in this experiment. The mean DNA content on day 0 post injection was set as 1.0, and that on day 15 post injection was 7.3. Unpaired t-test was used for comparison (*p* < 0.05). Data are from three biological replicates.

**S1 Table. Days confirming infection by administration of sporozoite of male and female-injected mosquito**. At 28 d post injection, 2000–10000 *P. berghei* sporozoites from whole bodies or salivary glands of the injected males and females, respectively, were administered into the tail vein of mice. Seven injected females were blood-fed on mice, and the infection in the mice was checked every day using a blood smear test. I.V.: administration into the tail vein.

